# Identification of leukemia-associated immunophenotypes by database-guided flow cytometry provides a highly sensitive and reproducible strategy for the study of measurable residual disease in acute myeloblastic leukemia

**DOI:** 10.1101/2022.07.12.499672

**Authors:** P Pinero, M Morillas, N Gutiérrez, E Barragán, E Such, J Breña, C Gil, C García, C Botella, JM Navajas, P Zapater, P Montesinos, A Sempere, F Tarín

## Abstract

**Background:** Multiparametric Flow Cytometry (MFC) is an essential tool to study the involved cell lineages, the aberrant differentiation/maturation patterns and the expression of aberrant antigens in acute myeloid leukemia (AML). The characterization of leukemia-associated immunophenotypes (LAIPs) at the moment of diagnosis is critical to establish reproducible strategies for the study of measurable residual disease using MFC (MFC-MRD).

**Methods:** In this study, we identified and characterized LAIPs by comparing the leukemic populations of 145 AML patients, using the EuroFlow AML/ MDS MFC panel, with 6 databases of normal myeloid progenitors (MPCs). Principal component analysis was used to identify and characterize the LAIPs, which were then used to generate individual profiles for MFC-MRD monitoring. Furthermore, we investigated the relationship between the expression patterns of LAIPs and the different subtypes of AML.

The MFC-MRD study was performed by identifying residual AML populations that matched with the LAIPs at diagnosis. To further validate this approach, the presence of MRD was also assessed by qPCR (qPCR-MRD). Finally, we studied the association between MFC-MRD and progression-free survival (PFS).

**Results:** The strategy used in this study allowed us to describe more than 300 different LAIPs and facilitated the association of specific phenotypes with certain subtypes of AML. The MFC-MRD monitoring based on LAIPs with good/strong specificity was applicable to virtually all patients and showed a good correlation with qPCR-MRD and PFS.

**Conclusions:** The described methodology provides an objective method to identify and characterize LAIPs. Furthermore, it provides a theoretical basis to develop highly sensitive MFC-MRD strategies.

## INTRODUCTION

The evaluation of measurable residual disease (MRD) is an essential decision-making tool in acute myeloblastic leukemia (AML) [1-5]. MRD monitoring, regardless of the technique employed, is used as an independent prognostic indicator for treatment planning [4, 6, 7]. The currently applied methods for the analysis of MRD are multiparameter flow cytometry (MFC-MRD) and real-time quantitative polymerase chain reaction (qPCR-MRD) [8-10].

Two separate approaches have been used to evaluate MFC-MRD: (1) the identification of leukemia associated immunophenotypes (LAIPs)-based strategy and (2) the study of different from normal maturation patterns (DfN)-based strategy [8]. To achieve fully reproducible results, both approaches require automated analysis strategies based on standardized procedures.

LAIPs consist in the aberrant expression of CD markers in leukemic cells (overexpression, lack or underexpression, asynchronisms and cross-lineage markers) [4, 8, 11-14]. The identification of LAIPs is the most frequently used strategy to study MFC-MRD in AML but has been exposed to great difficulties due to the application of non-standardized protocols, the limited information about their sensitivity and specificity, or the possible interference with normal minor populations [4, 11, 12, 14]. Although several harmonization studies have improved the concordance between different centers, standardized approaches including the use of automated analysis tools are the best way to obtain reproducible results [8, 15].

In order to minimize the subjectivity during the analysis of multiparametric data in AML, we designed a prospective study in which we used automated tools to compare leukemic populations with databases of normal immature myeloid precursors (MPs) in standardized 8-color AML experiments. Our objectives were to characterize LAIPs to identify which markers contribute to the optimal discrimination of AML cells, to investigate the association between LAIPs and different AML subtypes, and finally, to propose a MFC-MRD analysis strategy based on automated tools that can be shared by different researchers.

## METHODS AND STUDY DESIGN

### General study strategy

The general strategy used for this study is summarized in figure 1. First, we created six different databases from FCS files of normal bone marrow (BM) samples. Then, FCS files from BM samples of newly diagnosed AML patients were compared with each database. LAIPs were characterized by deviations in antigenic expressions, using multidimensional analysis strategies. Once the FCS files were analyzed, a specific follow-up profile including a reference image of the LAIP was created and stored. These profiles were later used to detect residual leukemic populations in samples from patients with complete remission (CR), as well as to establish the specificity of LAIPs in post-chemotherapy samples from non-AML patients.

**Figure 1.**
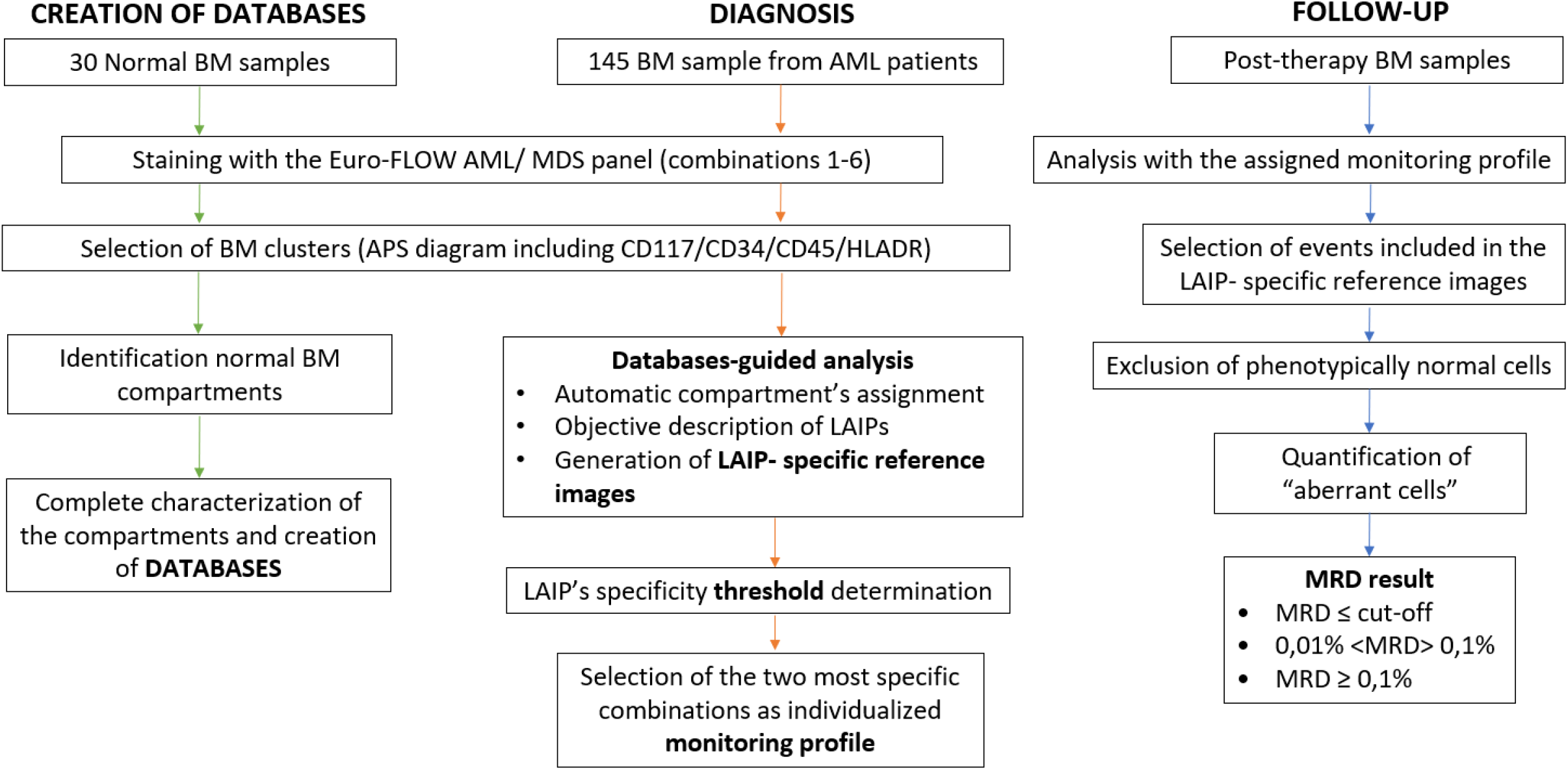
Flow diagram of the study.

### Patients and samples

BM samples from 145 AML patients diagnosed at the Hematology Unit of the General University Hospital of Alicante were included in the study of LAIPs. In addition, we collected 30 bone marrow samples from healthy donors obtained during orthopedic surgery procedures and 30 post-chemotherapy regenerated BM samples from non-AML patients (15 from multiple myeloma autologous transplant patients and 15 from post-induction acute lymphoblastic leukemia patients).

Sample processing was performed following standardized procedures validated for both cytometry and molecular studies [8, 15]. Patients were diagnosed and classified according to the revised 2016 World Health Organization criteria [16]. Clinical characteristics of patients are summarized in table 1.

**Table 1.** Characteristics of patients

| **CLINICAL CHARACTERISTICS (n=145)** |  |
| --- | --- |
| Sex (Male/Female) | 145 (78/69) |
| Mean age (range) | 61.9(18-92) |
| Leucocytes x 10<sup>9</sup>(range) | 26.3(0.3-323) |
| Hemoglobin (g/L) (range) | 93.5(35.3-143.3) |
| Platelets x 10<sup>9</sup>(range) | 44.9 (2-402) |
| % Bone marrow blasts (range) | 55.7 (11-99) |
| **AML SUBTYPES (WHO classification)** |  |
| **AML with recurrent genetic abnormalites** | **88 (60.7%)** |
| t(8;21) RUNX1-RUNX1T1 | 10 (6.9%) |
| Inv16 CBFB-MYH11 | 13 (8.9%) |
| t(15;17) PML-RARα | 11 (7.6%) |
| t(9;11) KMT2A-MLLT3 | 9 (6.2%) |
| t(6;9) DEK-NUP214 | 2 (1.4%) |
| Inv3 GATA2 | 1 (0.7%) |
| Mutated NPM1 | 32 (22.1%) |
| Bialelic mutation of CEBPA | 5 (3.4%) |
| Mutated RUNX1 | 4 (2.8%) |
| t(9;22) BCR-ABL | 1 (0.7%) |
| **AML with myelodisplasia-related changes** | **15 (10.3%)** |
| **Therapy-related neoplasms** | **5 (3.4%)** |
| **AML, NOS** | **37 (25.5%)** |
| AML with minimal differentiation | 7 (4.8%) |
| AML without maturation | 9 (6.2%) |
| AML with maturation | 6 (4.1%) |
| Acute monoblastic and monocytic leukemia | 12 (8.3%) |
| Acute erithroid leukemia | 2 (1.4%) |
| Acute megakaryoblastic leukemia | 1 (0.7%) |
| **Karyotype** |  |
| Normal | 28 (19.3%) |
| Recurrent | 46 (31.7%) |
| Isolated | 18 (12.4%) |
| complex | 15 (10.3%) |
| not valuable | 28 (19.3%) |
| **Other molecular findings** |  |
| FLT3 | 34 (22.8%) |
| DNMT3A | 17 (11.7%) |
| IDH1/IDH2 | 13 (9.1%) |
| N-RAS | 12 (8.3%) |
| ASXL1 | 7 (4.8%) |
| SRFS2 | 11 (4.1%) |
| Others | 37 (22.1%) |
| Non determined | 26 (17.9%) |

After diagnosis, 106 patients received at least one induction cycle following the standard “3+7” first-line regimen (Idarubicin 6-10 mg / m2 day 1-3 and Ara-C 100-150 mg / m2 day 1-7). The remaining patients (39) received other treatments and were excluded for MFC-MRD monitoring. 73 of the 106 patients (69.5%) achieved CR after the first “3+7” cycle and were eligible for successive MRD monitoring (supplementary table 1). The quality of normal and MRD samples was assessed by performing an automatic count of nucleated cells as well as red blood cell percentage, CD34+ cells, CD117+ myeloid precursors, B cell precursors, plasma cells and mast cells.

This study has been approved by the Ethics Committee of our Hospital and all participants signed an informed consent, in accordance with the Declaration of Helsinki.

### Flow cytometry

Eight-color MFC was performed on BM samples within 24 hours after the extraction. All experiments, both at the moment of diagnosis and during follow-up, were performed following the EuroFlow Standard Operating Procedures for instrument setup, fluorescence compensation and sample preparation [17]. Samples were stained with the combinations 1 to 6 of the EuroFlow standardized AML/MDS panel [18].

The MFC-MRD studies were performed using 300 μL of BM specimens from patients with CR (100 μL/tube), and the two most informative combinations (according to the specificity of the identified LAIP) were reproduced during the follow-up. The acquisition was performed on a FACS-CANTO II flow cytometer with DIVA software (BD Biosciences, CA). FCS files (comprising a minimum of 1.000.000 cells after exclusion of debris, doublets, and erythrocytes) were analyzed using Infinicyt Flow Cytometry Software (Cytognos, Salamanca, Spain, V 2.0).

### Normal Databases

We created six normal databases using FCS files from BM samples from healthy donors. Each database contained the characteristics of the distinct normal myeloid compartments defined by the antibodies included in the panel. Each compartment shows a particular immunophenotype that allows the comparison between normal and aberrant myeloid progenitors at different maturation stages. The strategy used to select the normal compartments, as well as their phenotypic characteristics, is detailed in figure 2. LAIPs characterization and description and MFC-MRD analysis was made by two independent researchers.

**Figure 2.**
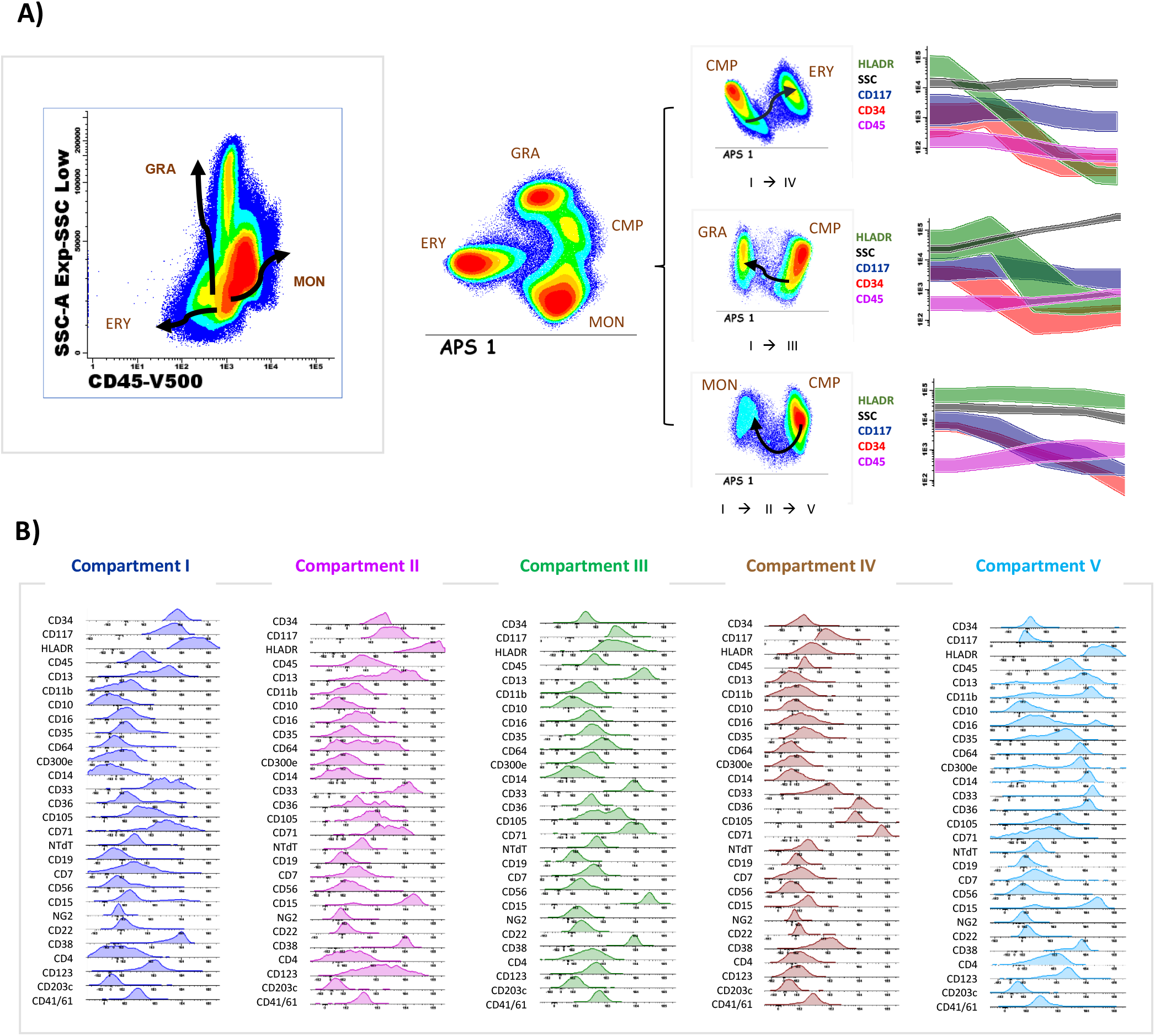
**A)** Identification of 5 different maturational compartments attending to the expression of the backbone markers (CD45/CD34/CD117/HLADR) and SSC in an APS graph. Changes in the intensity of the backbone markers throughout consecutive maturational compartments. The common myeloid progenitor (CMP) is represented by compartment I. The monocytic differentiation (MON) occurs from compartment I to compartment II and finally compartment V. The granulocytic differentiation (GRA) arises from compartment I towards compartment III. Erythroid differentiation (ERY) derives from the common myeloid progenitor to compartment IV. **B)** Complete immunophenotype of the identified compartments (I to V) using the Euroflow standardized AML/SMD panel.

### LAIPs characterization and description and MFC-MRD analysis

At diagnosis, the leukemic populations were selected using a common gating strategy based on the expression of the four backbone markers and SSC in a multidimensional diagram (fig. 3-A). Subsequently, the leukemic population was automatically classified according to the closest maturation compartment (figure 3-B).

**Figure 3.**
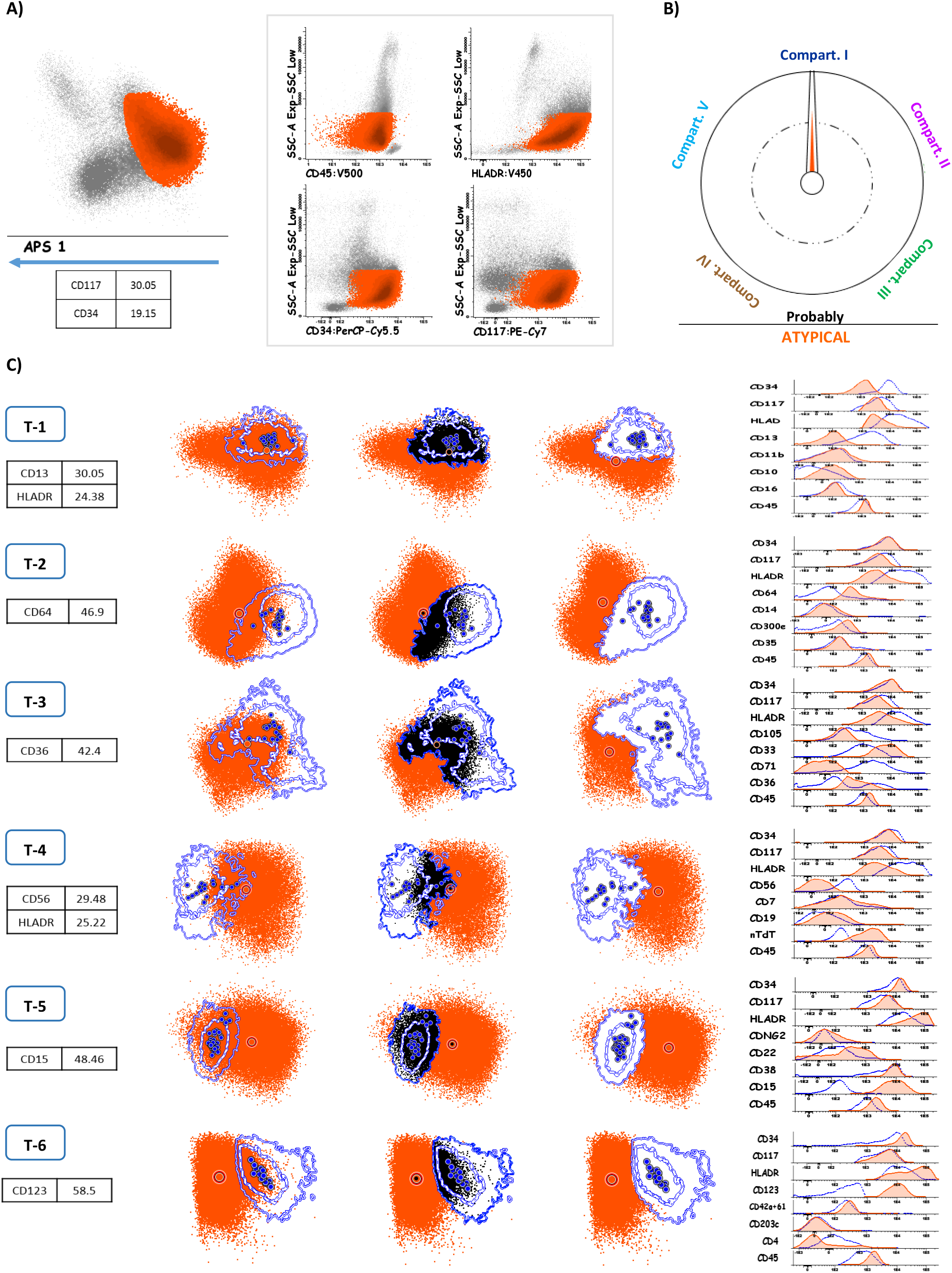
Phases of the analytical process at AML diagnosis. **A)** Identification and selection of the leukemic population attending to the backbone markers by using an APS graph. **B)** The compass tool included in the normality databases automatically classifies the population, the orange arrow points to the assigned compartment. **C)** Comparison of the leukemic cluster with the assigned normal compartment. The overlapping events, indistinguishable from the normal phenotype, are excluded from the analysis. The description of the LAIP is based on the main differences with respect to the indicated compartment (as shown in the histograms).

The populations were compared with the indicated compartment and the main phenotypic differences were identified. LAIPs were defined as cell populations separated from all the compartments represented in the databases and were described indicating only the most informative markers contributing to the separation from the compartment to which it was assigned (fig 3-C).

### Generation of individualized monitoring profiles

To generate individualized monitoring strategies, we created specific reference images of each aberrant population. Leukemic events that, with the antibody combinations tested, overlap with the normal compartments (indistinguishable from normal) were excluded. Consequently, only events located in empty spaces were stored in the reference images (figure 4 A-B), which enabled us to have a specific strategy for the identification and quantification of MFC-MRD populations (figure 4C).

**Figure 4.**
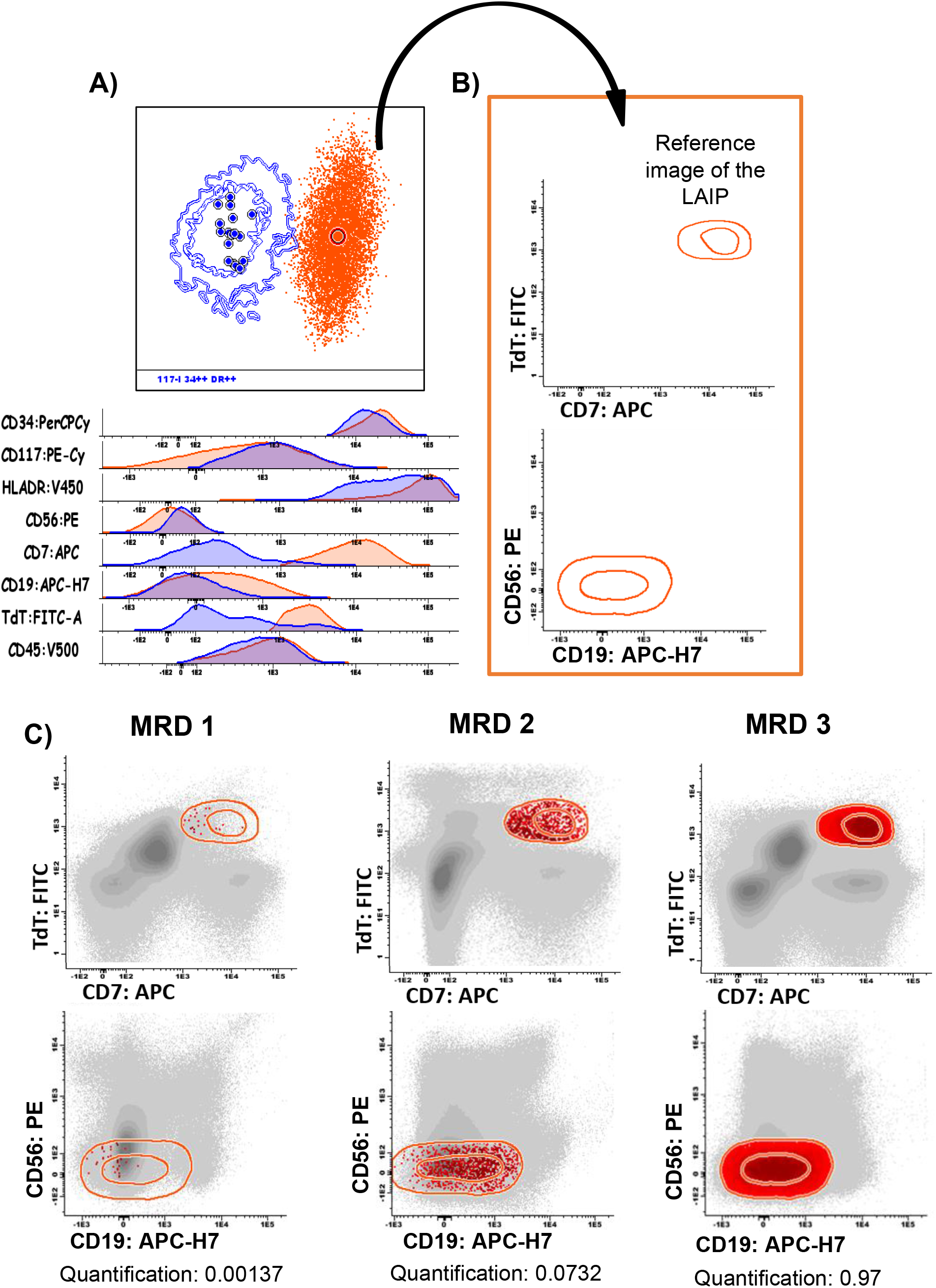
Creation of the reference images for the successive MRD monitoring. **A)** The leukemic population at diagnosis (orange) was compared with its theoretical normal counterpart (blue) and the LAIP was defined as follows: compartment I-CD7^hi^, TdT^hi^, CD19^hi^, CD56^hi^. **B)** A reference image of the leukemic cluster was automatically created to investigate the LAIP’s cut-off in normal/regenerative samples. This case showed a high sensitivity LAIP (individual threshold of 0.00154) at diagnosis. **C)** Results of three successive MRD evaluations. MRD-1= negative (<threshold), MRD-2= positive* (<0.1%, but> threshold) and MRD-3= positive (>0.1%)

### Specificity of LAIPs in regenerative bone marrow samples

With the aim of determining the specificity of LAIPs, we evaluated their presence in a fusion of 30 FCS files of post-chemotherapy regenerated BM samples. We used the reference image of each individual LAIP to detect the events included therein (+2 SD). The quantification of the LAIPs in these merged files provided useful data to establish an individual LAIP-specific threshold in subsequent MFC-MRD analysis.

### MFC-MRD analysis

The assessment of MFC-MRD study was performed in samples from patients with CR using the previously stored follow-up profiles. A positive MFC-MRD was established when a homogeneous cluster was detected expressing an immunophenotypic profile equivalent to the original LAIP and exceeding the LAIP-specific threshold. This population was quantified once the erythroid population was excluded.

### PCR-based MRD analysis

The qPCR analyses were performed in bone marrow samples according to the European Leukemia Net [8] in all CBF-MYH11, RUNX1-RUNX1T1 and most of NPM1+ patients. The follow-up by qPCR-MRD was performed at the same time-points as MFC-MRD.

### Next-generation sequencing

Next generation sequencing (NGS) analyses were performed at diagnosis. For this, the 40 genes included in the Oncomine myeloid research panel (Thermo Fisher Scientific) were evaluated.

### Statistical methods

Continuous variables are reported as means ±standard deviations (SD) and ranges. Categorical variables are given as frequency or percentages. To define the individual thresholds of LAIPs we established the cut-off in the upper value of the 95%-confidence interval, according to the Poisson approximation. A hierarchical clustering analysis was performed to identify relevant clusters based on immunophenotypic characteristics. Chi-square test was used to determine the association between categorical variables. Receiver Operating Characteristic (ROC) curves were used to study the discriminating ability of MFC-MRD, using qPCR-MRD results as reference method. Progression-free survival (PFS) curves were constructed using the Kaplan-Meier method and compared with the log-rank test. All analyses were performed with the SPSS statistical software, version 15.0 (IBM, Armonk, New York).

## RESULTS

Our strategy allowed us to identify 310 different LAIPs that represented variable percentages of the leukemic population (LAIP size, mean 49.9%, range 1.18%-99.9%). The average number of LAIPs per case was 3.73. The mean specificity of the LAIPs was 0.0845 (range: <0.001%-3.11%). No significant correlation was demonstrated between the percentage of cells carrying the LAIP and its specificity (R^2^= -0.11; p>0.1). LAIPs were classified in 3 “specificity categories” as described by Rossi and colleagues [4]. As a result, 31% of the LAIPs were classified as strong LAIPs (LAIP specific threshold <0.01%), 48% as good LAIPs (LAIP specific threshold 0.01-0.1%) and 21% as weak LAIPs (LAIP specific threshold>0.1%). Following this classification, 96.7% of the patients had at least 2 LAIPs of strong or good specificity. The number of LAIPs was similar in all the EuroFlow combinations, although the combinations 1, 3, 5 and 6 showed stronger LAIPs than combinations 2 and 4. (supplementary table 2).

### Classification and characteristics of LAIPs

Using the strategy described above, we were able to classify leukemic clusters according to the closest maturation compartment. Using this strategy, 51.7% of cases were classified into compartment I, 10.3% into compartment II, 25.5% into compartment III, 2.1% into compartment IV and 10.3% into compartment V.

LAIPs showed different characteristics depending on the normal compartment to which they were referred to. In the compartment I, alterations consisted mainly in combinations of two markers per tube (including changes in the intensity, asynchronous and cross-lineage expression of several antigens) and mostly corresponded to the categories of strong/good LAIPs (table 2). As a single marker, the overexpression of CD123 was the most frequent abnormality detected in the combination 6 (Figure 5).

**Table 2.**
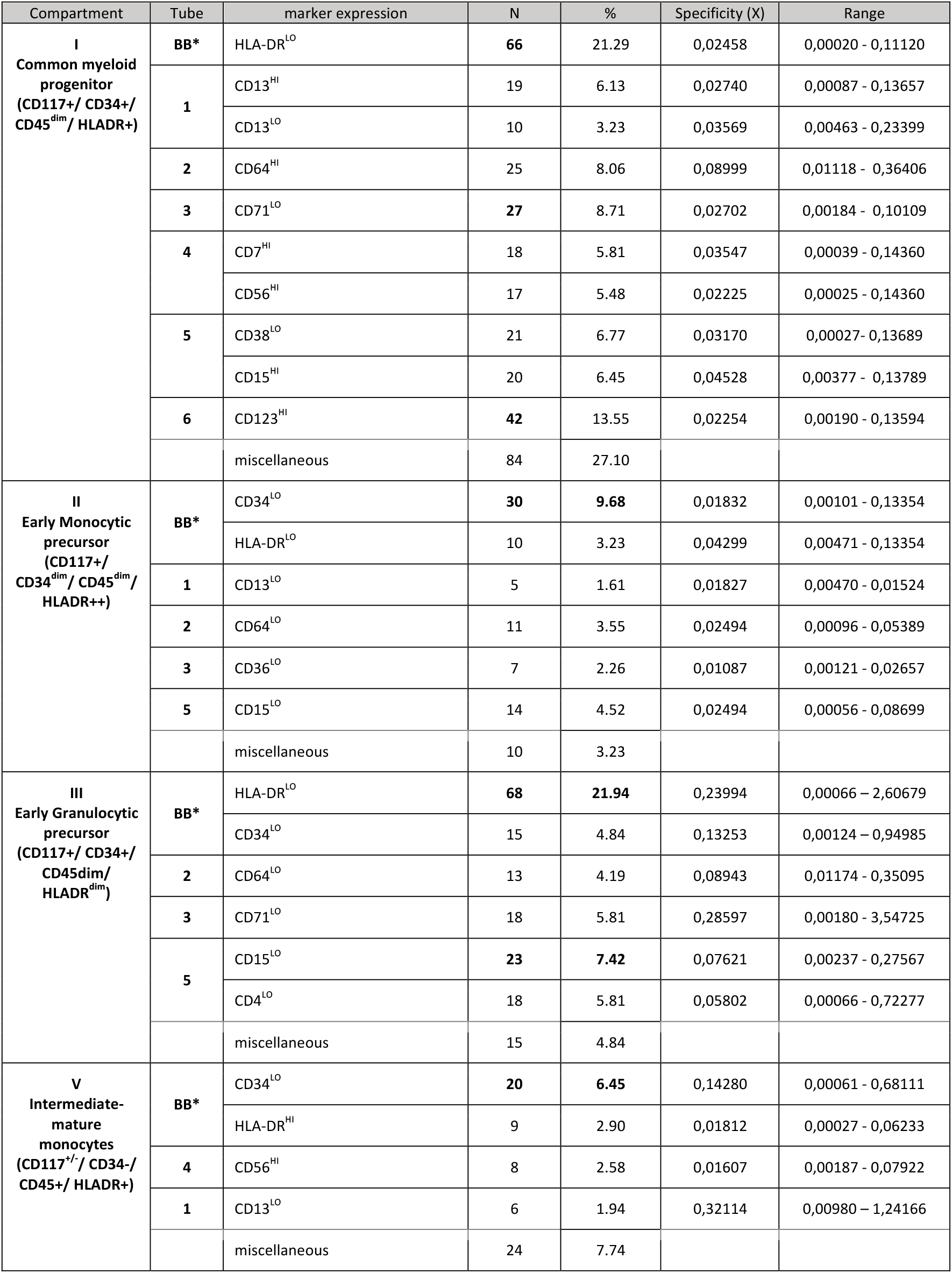
Distribution of LAIPs in the different compartments

**Figure 5.**
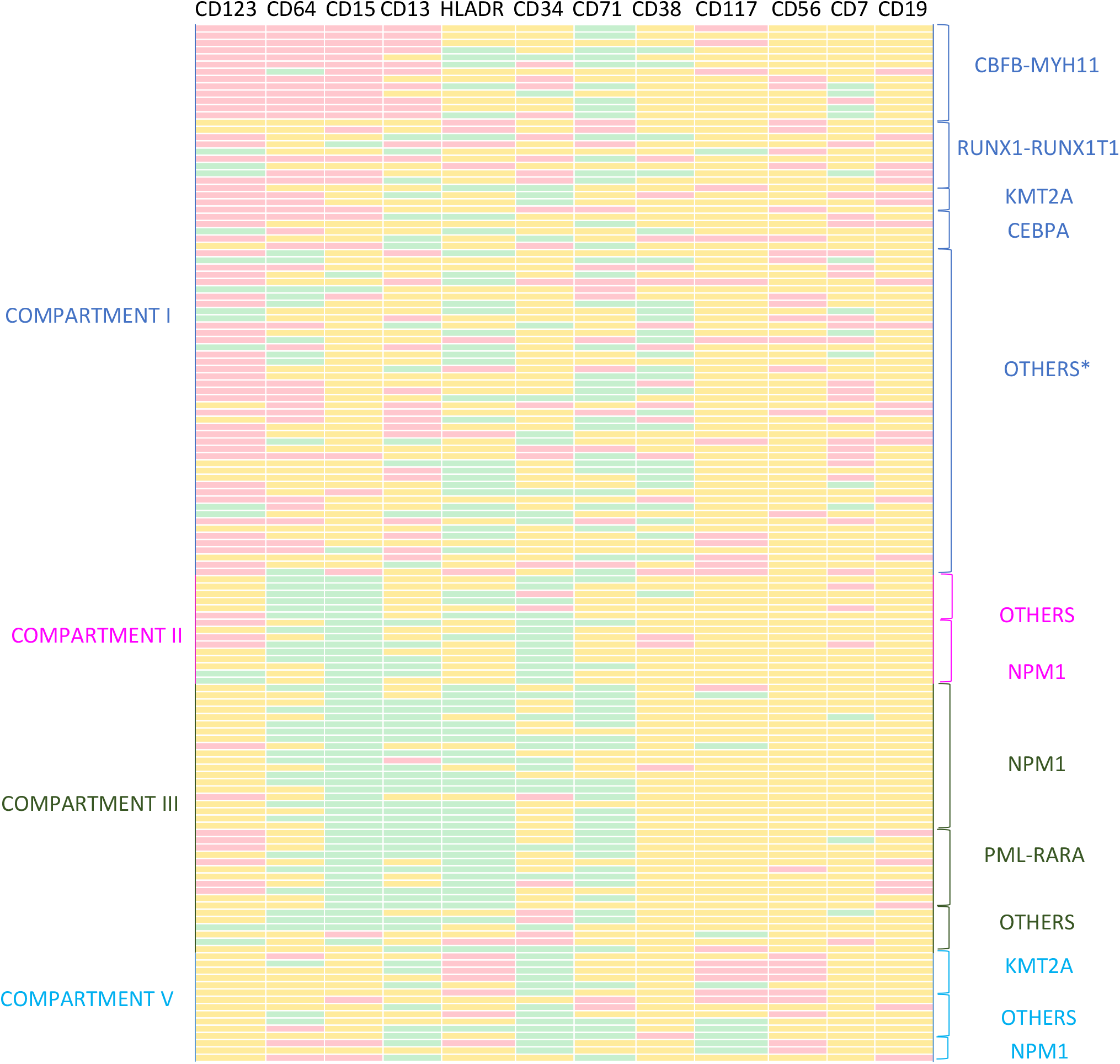
Immunophenotypic profiles in the different subtypes of the AML-WHO classification. The intensity of expression for the mainly altered antigens (represented in columns) is represented in yellow, red or green depending on whether it is a normal expression, overexpression or under-expression respectively. * In the compartment 1, the group “others” included RUNX1 mutated, BCR-ABL, DEKNUP, NPM1, MDS-related changes, therapy-related myeloid neoplasms and NOS cases. In the remaining compartments the group “others” included only NOS and MDS-related changes cases.

In compartment II, asynchronisms and changes in the intensity of several antigens were the predominant alterations, whereas cross-lineage expressions were only rarely found. Most of the LAIPs showed a decreased expression of CD34 in combination with HLADR^low^, CD13^low,^ CD64^low^, CD36^low^ and/or CD15^low^ (Figure 5). LAIPs mostly corresponded to strong/good LAIPs in a similar way to those of compartment I (Table 2).

In compartment III, the most frequent alterations included different combinations of HLADR^low^ with CD34^low^, CD64^low^, CD71^low^, CD15^low^ or CD4^low^ (Figure 5). In this group, the specificity of LAIPs was variable, with a significant prevalence of weak LAIPs (Table 2).

There were only 3 cases assigned to compartment IV (2 AML-M6, 1 AML-M7) so the results in this group were not included in the analysis.

In compartment V, CD34^low^, HLADR^hi^, CD13^lo^ and CD56^hi^ were the most frequently found aberrancies (Figure 5). Several weak LAIPs were found, usually involving heterogeneous expression of the previously mentioned markers. In this group, the high expression of HLADR or CD56 was characteristically associated with strong/good LAIPs (Table 2).

### LAIPs and AML world health organization (WHO) molecular subtypes

The results of cytogenetic and molecular studies were unknown at the time of MFC evaluation. However, the subsequent incorporation of data showed that the different AML-WHO subtypes were strongly associated with certain maturational compartments and immunophenotypic profiles (figure 5). Compartment I included all AML CBFβ-MYH11 and RUNX1-RUNX1T1, as well as less frequent variants of AML (biallelic mutations of CEBPA), mutated RUNX1, BCR-ABL and DEK-NUP214 cases). Three KMTA-MLLT3 cases were also included in this compartment. Regarding the relationship between specific immunophenotypic characteristics and AML subtypes, we identified a characteristic phenotype in CBFβ/MYH11 AML cases (CD123^hi^/ CD64^hi^/ CD15^hi^/CD13^hi^). RUNX1/RUNX1T1 cases showed a variable immunophenotype with frequent alterations in CD13, CD34, HLADR, CD15, CD123 and CD71, as well as expression of cross lineage antigens (mainly CD56 and/or CD19). The few cases of KMT2A AML showed a particular phenotype characterized by a lower expression of CD34, frequently accompanied by CD64^hi^ and CD123^hi^. Most of the AML cases without recurrent genetic abnormalities [including AML with MDS related changes, not otherwise specified (NOS) and therapy-related cases] showed a similar phenotype with changes in CD38 (usually CD38^lo^) and characteristics of early progenitors. Alterations in the expression of CD71, HLA-DR, CD123 and CD7 were also observed.

Compartment II included some mutated NPM1 cases with evidence of monocytic differentiation. Their characteristic immunophenotype included CD15^lo^, CD34^lo^ and often CD64^lo^, but still indistinguishable from the NOS cases included in this compartment.

Compartment III included all PML-RARα and mutated NPM1 cases that evidenced granulocytic differentiation. Both AML subtypes showed a similar phenotype (CD15^lo^, HLADR^lo^, CD13^lo^ and CD71^lo^) although some PML-RAR cases showed overexpression in CD19. On the other hand, we did not identify a particular immunophenotype associated with FLT3 alterations.

Compartment V included CD34-AMLs with evident monocytic maturation. Most KMT2A-AML cases were included in this compartment and presented a recognizable phenotype (CD117^hi^, HLADR^hi^, CD56^hi^) that allowed us to differentiate them from remaining monocytic variants. Some NPM1+ AML monocytic variants were also included in this group and showed a characteristic CD300e^hi^ phenotype.

### MFC-MRD study and progression-free survival

We studied the association between MFC-MRD and PFS in 75 patients that reached CR after the first induction cycle. We tracked the two LAIPs that showed highest specificity in each patient. Interestingly, 97.4% of AML cases which did not meet the requirements for molecular monitoring showed at least two good LAIPs and 53.2% showed at least two strong LAIPs. The most repeated follow-up combinations were, in this order: 1+6 (33%), 1+3 (19%), 1+4 (15%), 2+4 (12%), 1+5 (12%) and 2+6 (5%) and others (4%).

Patients were stratified into three categories, MFC-MRD>0.1%, MFC-MRD 0.1-0.01% and MFC-MRD< LAIP specific threshold as in previous studies [14]. Patients with MFC-MRD<0.1% showed an estimated PFS of 39.2 ± 3.3 months (95% CI: 31.5-46.9) (Figure 6A), that was significantly lower than both MFC-MRD 0.1-0.01% (50.5 ± 4.3 months; 95%CI: 41.8-59.0) and MFC-MRD< LAIP specific threshold (mean 53.7 ± 4.6 months; CI95% 44.7-62.5) (Figure 6B-C). Interestingly, the last two groups showed almost the same PFS (figure 6B-C).

**Figure 6.**
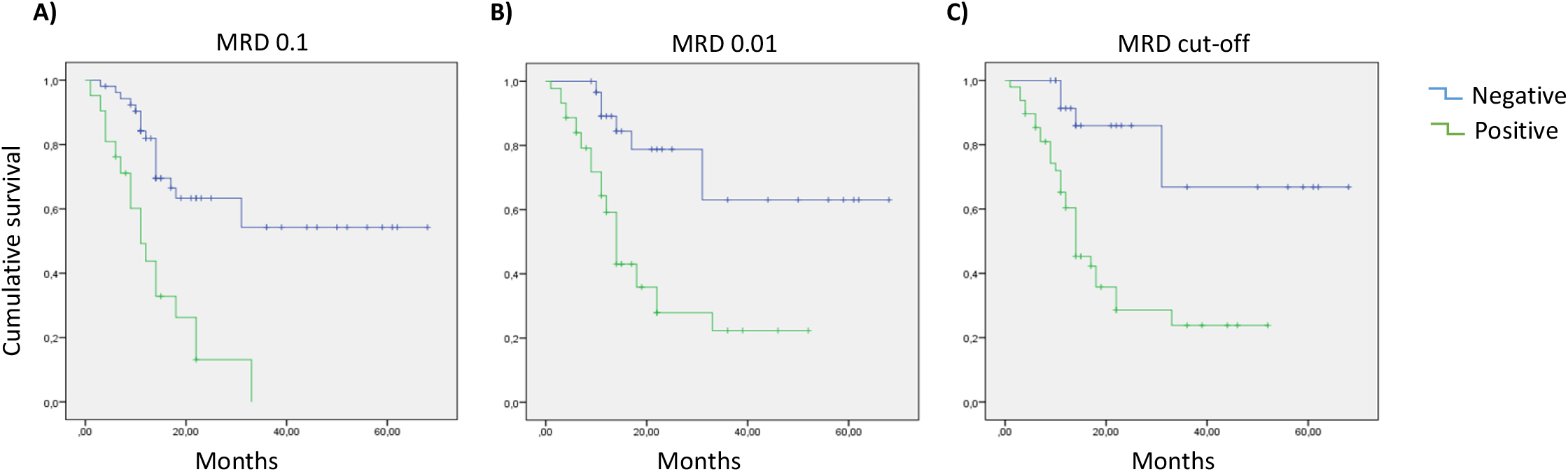
Kaplan-Meier survival curves for AML patients using 3 different cut- off for the evaluation of MRD in remission samples after first induction cycle. **A)** Time to relapse in AML patients using a MRD threshold of 0.1. (PFS= 39.18 ± 3.3 months; p=0.001). **B)** Time to relapse in AML patients using a MRD threshold of 0.01. (PFS= 50.5 ± 4.3 months; p<0.001). **C)** Time to relapse in AML patients using individual MRD threshold. (PFS= 53.7 ± 4.6 months; p<0.001)

### MFC-MRD and qPCR-MRD

To study the correlation between MFC-MRD and qPCR-MRD, we compared the results obtained by both methods in 203 successive CR samples subsidiaries of molecular monitoring (CBFB/MYH11, RUNX1/RUNX1T1 and mutated NPM1 variants). The MRD results included 55 (27.1%) qPCR positive cases and 148 (72.9%) qPCR negative cases. Our results showed that 46 samples (22.6%) were MFC-MRD+ (>individual cut-off) and qPCR-MRD+, whereas 138 samples (67.9%) were negative by both methods. Discordant results were observed in 19 samples (9.4%): 9 samples (4.4%) were MFC-MRD-but qPCR-MRD+ and 10 samples (4.9%) were MFC-MRD+ but qPCR-MRD-. Thereby, we confirmed a good concordance of these methods (Kappa coefficient of 0.692; 95%CI: 0.580-0.797).

Furthermore, when the samples were classified as positive or negative attending only to the qPCR results, levels of MFC-MRD were much higher in the qPCR+ group than those in the qPCR- (t test, p< 0.001). We also determined that MFC-MRD cut-off value of 0.0087% improves the specificity (80.6%) without impairing sensibility compared with standard cut-off of 0.1% for MFC-MRD evaluations.

## DISCUSSION

In this study we have developed a database-guided strategy to detect and characterize LAIPs in newly diagnosed AML patients. Previous studies have developed different automated supervised strategies, based on standardized EuroFlow protocols, to identify the leukemic lineage and correlate the phenotype with some molecular findings. [18, 19]. Our strategy goes further allowing the identification of LAIPs even if they are expressed only in a small part of the leukemic population, which are otherwise impossible to detect by conventional methods. Thus, our findings confirm previous observations evidencing that conventional strategies could be significantly improved [4, 20-22]

One of the greatest challenges for MFC-MRD assessment is the presence of cell populations expressing the identified LAIPs in healthy or regenerating bone marrow samples [8, 23]. Our results confirm these observations even using individual monitoring profiles and database guided analysis for the exclusion of phenotypically normal cells. The presence of this normal background limits MFC-MRD studies, although the addition of more fluorochromes and the implementation of automatic procedures may probably reduce its importance.

To overcome this difficulty, different studies have stratified LAIPs into specificity categories, according to their phenotypic characteristics, assigning a specific threshold equivalent to their count in regenerative non AML samples [4, 24, 25]. Although we have followed this methodology, it should be noted that the same LAIP has shown different specificities depending on the combination of antibodies used. Furthermore, the contribution of apparently less informative markers can significantly modify the specificity of LAIPs. Therefore, we strongly recommend to describe LAIPs according to the full combination of markers used, select the most specific LAIPs for MRD studies and to reproduce the same combinations for diagnosis and follow-up [8, 15]

In this study, we propose a systematic classification of LAIPs based on the maturational stage in which they are found, providing a simplified nomenclature to facilitate their description and characterization by specifying the combination of antibodies that allows their identification. In addition, our results confirmed that the most specific LAIPs are found in the most immature compartments, as has been previously described [4, 8, 23-25], highlighting the importance of analyzing the most immature compartments. Some exceptions were observed in monocytic leukemias with high expression of CD56 or HLA-DR, although the stability of both markers could be affected in successive relapses, which limits their value for future MFC-MRD evaluations [26].

It is well known that certain AMLs display a particular immunophenotype that correlates with the WHO molecular subtypes and certain genetic abnormalities [27-29]. Our study has allowed us to correlate LAIPs with specific WHO AML variants. Particularly, CBF/MYH11+ AML progenitors showed an almost exclusive immunophenotypic signature, as well as several characteristic LAIPs that were found in other molecular subtypes such as KMT2A, PML-RARa and NPM1. In addition, most of the cases without recurrent genetic abnormalities (including AML with MDS related changes, NOS and therapy-related cases) showed a similar phenotype, usually CD34+ CD38^lo^ that has been related to an immature phenotype and properties of leukemic stem cells[30-32].

These results, beyond their possible usefulness to refine the diagnosis, support the validity of the strategy to characterize LAIPs, since they bring consistent data related to the biological characteristics of each AML subtype. To investigate the possible application of our strategy, we have studied the association between MFC-MRD results with progression-free survival. Our results suggested that an optimal cut-off for MFC-MRD would be 0.01%, since it closely correlates with the risk of relapse and discriminates better than 0.1% between patients with different PFS. Previous data have already advised that the currently applied cut-off (0.1%) is not accurate enough to identify patients with a good prognosis [8].

An important observation is the high frequency of strong LAIPs in patients who are not subsidiary to qPCR monitoring, revealing that MFC-MRD 0.01% cut-off would be applicable to most of these patients and leaving only a small proportion of cases that should be monitored with alternative strategies. Recent experiences have already suggested that NGS and MFC are synergistic and could improve the results of the MRD detection in a combined strategy [33, 34].

We have monitored patients by qPCR and MFC and we have demonstrated a high concordance between successive MFC-MRD and qPCR-MRD results. Based on the qualitative qPCR result (positive/ negative) we have confirmed that a generic cut-off value close to 0.01% also provides an optimal sensitivity and specificity for MFC-MRD evaluations.

A theoretical limitation of the proposed strategy would be the emergence of clones with phenotypic shifts [13, 35]. Though, our results showed that a fraction of the LAIPs remain stable when the molecular disease persists and is associated with a high risk of relapse. Considering that, we highlight the importance of the detection and monitoring of, at least, two LAIPs in every patient. Further studies on larger panels and alternative approaches such as the DfN strategy would be necessary to avoid the dependency on the original LAIPs. But, in any case, automated analysis tools and comparison with databases seem to be essential for future diagnostic strategies.

In conclusion, our work provides an objective strategy for the characterization of LAIPs and the monitoring of MFC-MRD in standardized eight-color MFC experiments. To our knowledge this is the largest study investigating the relevance of the diagnostic standardized EuroFlow AML/MDS panel on MRD evaluations so far. Additionally, given that databases could be shared in collaborative projects, our strategy represents a promising onset to reach a consensus in MFC-MDR studies.

