## Supplementary material for "Identification of leukemia-associated immunophenotypes by database-guided flow cytometry provides a highly sensitive and reproducible strategy for the study of measurable residual disease in acute myeloblastic leukemia": S2

**Supplementary table 2.**

| | **Specificity** mean (range) | **<0.01%** | **≥0.01-0.1%** | **>0.1** |
|---|---|---|---|---|
| **T1** | 0,0845 (0,0002636- 1,24165) | 28 (32%) | 43 (49%) | 16 (18%) |
| **T2** | 0,1035 (0,0009616- 2,6067938) | 17 (16%) | 49 (46%) | 34 (32%) |
| **T3** | 0,0843 (0,00020497- 3,547254) | 43 (42%) | 43 (42%) | 15 (15%) |
| **T4** | 0,1104 (0,000247- 2,54468) | 27 (30%) | 45 (50%) | 18 (20%) |
| **T5** | 0,0814 (0,0002615- 2,215884) | 30 (30%) | 48 (48%) | 21 (21%) |
| **T6** | 0,0481 (0,00066- 0,8041249) | 39 (39%) | 50 (50%) | 11 (11%) |

Number of different LAIPs detected in the 6 first combinations of the AML/MDS panel. LAIPs were classified in 3 categories of specificity: strong, good and weak (<0.01%, ≥0.01-0.1% and >0.1). General specificity (mean + range) of LAIPs in each single combination.
*T1-6: tube 1-6
