## Supplementary material for "Identification of leukemia-associated immunophenotypes by database-guided flow cytometry provides a highly sensitive and reproducible strategy for the study of measurable residual disease in acute myeloblastic leukemia": S1

**Supplementary table 1.**

| | |
|---|---|
| **Patients eligible for MRD study (CR1 after 3+7 regimen)** | **75** |
| **Post-remission therapy** | |
| Chemotherapy alone | 54 |
| ATSP | 10 |
| Allo-TMO | 11 |
| **MRD controls** | |
| MRD-MFC post cycle 1 | 75 |
| MRD-MFC and qPCR post cycle 1 | 42 |
| Sucessive MRD-MFC and qPCR monitoring | 161 |
| MRD-MFC and qPCR post cycle 2 | 39 |
| MRD-MFC and qPCR post cycle 3 o + | 69 |
| **Patients non eligible for MRD study** | **70** |
| **Induction Chemotherapy** | |
| 3+7 regimen (no CR) | 22 |
| ATRA based regimens (acute promyelocytic leukemia) | 11 |
| 5-azacytidine based regimens | 15 |
| Other treatments | 9 |
| Palliative care | 9 |
| Early deaths | 4 |
| **Post-remission therapy** | |
| Chemotherapy | 21 |
| ATSP | 5 |
| Allo-TMO | 6 |
